# Sharpened and mechanically durable carbon fiber electrode arrays for neural interfacing

**DOI:** 10.1101/2021.01.21.427697

**Authors:** Elissa J. Welle, Joshua E. Woods, Ahmad A. Jiman, Julianna M. Richie, Elizabeth C. Bottorff, Zhonghua Ouyang, John P. Seymour, Paras R. Patel, Tim M. Bruns, Cynthia A. Chestek

**Affiliations:** The Biomedical Engineering Department at the University of Michigan, Ann Arbor, MI 48109 USA; The Electrical Engineering and Computer Science Department at the University of Michigan, Ann Arbor, MI 48109 USA; The University of Michigan, Ann Arbor, MI 48109 USA. He is now with the Electrical and Computer Engineering Department at King AbdulAziz University, Jeddah, Saudi Arabia; The University of Michigan, Ann Arbor, MI 48109 USA. He is now with the Neurosurgery Department at the University of Texas, Health Science Center, Houston, TX 77030 USA

**Keywords:** carbon fiber, dorsal root ganglia, electrophysiology, motor cortex, neural electrode, peripheral nerve interfacing, vagus nerve

## Abstract

Bioelectric medicine treatments target disorders of the nervous system unresponsive to pharmacological methods. While current stimulation paradigms effectively treat many disorders, the underlying mechanisms are relatively unknown, and current neuroscience recording electrodes are often limited in their specificity to gross averages across many neurons or axons. Here, we develop a novel, durable carbon fiber electrode array adaptable to many neural structures for precise neural recording. Carbon fibers were sharpened using a reproducible blowtorch method that uses the reflection of fibers against the surface of a water bath. The arrays were developed by partially embedding carbon fibers in medical-grade silicone to improve durability. We recorded acute spontaneous electrophysiology from the rat cervical vagus nerve (CVN), feline dorsal root ganglia (DRG), and rat brain. Blowtorching resulted in fibers of 72.3 ± 33.5 degree tip angle with 146.8 ± 17.7 μm exposed carbon. Silicone-embedded carbon fiber arrays were durable with 87.5% of fibers remaining after 50,000 passes. Observable neural clusters were recorded using sharpened carbon fiber electrodes from rat CVN (41.8 μV_pp_), feline DRG (101.1 μV_pp_), and rat brain (80.7 μV_pp_). Recordings from the feline DRG included physiologically-relevant signals from increased bladder pressure and cutaneous brushing. These results suggest that this carbon fiber array is a uniquely durable and adaptable neural recording device. In the future, this device may be useful as a bioelectric medicine tool for diagnosis and closed-loop neural control of therapeutic treatments and monitoring systems.

## I. Introduction

BIOELECTRIC medicine therapies use electrical stimulation to treat disorders of the nervous system [1]. Well-established uses of bioelectric medicines include sacral nerve stimulation for bladder diseases [2], cervical vagus nerve (CVN) stimulation for epilepsy [3], and deep brain stimulation for Parkinson’s disease [4]. More recently, CVN stimulation has been used to treat inflammation [5] and therapy-resistant depression [6]. In order to precisely modulate the function of target organs, devices incorporate unique electrode configurations, materials, stimulation patterns, and closed-loop control [7], [8]. Current electrode interfaces for neural stimulation are large extraneural leads [9] or nerve cuffs [10], [11]. These interfaces are generally effective for stimulation but are limited in selectivity, often leading to side effects [12]. Furthermore, these interfaces have minimal utility for monitoring neural signals, which requires specific, multi-channel recording of the nerve or target organ. Current open-loop bioelectric medicine therapies may increase in efficacy if recording devices delineate the mechanisms of organ control or obtain organ-state signals.

Towards that end, intraneural interfaces offer greater recording signal-to-noise ratio and stimulation selectivity than extraneural interfaces [13], [14]. Clinical intraneural interfaces include the Utah Slanted Electrode Array [15], [16] and the longitudinal and transverse intrafascicular electrode arrays [17]–[19]. The selective access to axons and fascicles enabled by conventional intraneural interfaces has facilitated clinical research of nerve-machine interfaces [20], which connect peripheral nerves to an external robotic device, allowing the nervous system to send motor commands to [21], and receive sensory input from the device [22]. However current interfaces have limited long-term viability due to electrode material failure [23], [24] and significant scarring [25], [26]. Additionally, their size is not appropriate for the small nerves of the autonomic nervous system, which provide primary organ innervation. An ideal interface with autonomic nerves would maximize recording specificity while minimizing scarring, maintain a high channel count of small-footprint electrodes, and contain durable, biocompatible electrode materials.

Some recent interfaces are nearing these design criteria. Novel electrode development often starts with cortical applications that may later undergo translation to peripheral target locations. An injectable mesh-style PDMS array with 10 μm-diameter elements prompted minimal histological scarring and maintained long-term recording in the brain [27], [28]. However, the injection method of insertion would be difficult to use in peripheral applications. Flexible carbon nanotube “yarn” electrodes recently demonstrated physiologically-relevant recordings in the rat CVN and glossopharyngeal nerves for up to 16 weeks with minimal scarring [29]. This suggests that ultra-small electrodes may be similarly biocompatible with nerves, as has been shown in the brain [30]–[32]. While capable of recording from awake animals, the fragile “yarn” material and manual insertion of each electrode with a shuttle or suture may limited the channel count of this design [33].

Neural interfaces constructed of rigid components at micron size scale may ease surgical implementation while still minimizing tissue response. A novel design uses carbon fiber electrodes, which have been primarily used for brain recordings [34]–[36], as a method to record from nerves [37]. The carbon fiber electrodes reported in Gillis et al., 2017 successfully recorded acute spontaneous spiking activity [37]. However, the majority of their recordings were evoked responses to electrical stimulation, which are not as indicative of chronic in-dwelling performance as smaller-amplitude spontaneous neural signals. Additionally, the architecture of their array would be difficult to translate from acute to chronic implantation. The carbon fibers’ rigid junction to the substrate provides a likely breakage point. An array with sub-cellular carbon fiber electrodes that is compatible with peripheral geometry and capable of spontaneous neural recording is needed for neural interfacing.

Here, we develop and demonstrate a carbon fiber array comprised of several novel components: sharpened carbon fibers (SCFs), silicone-embedded carbon fibers, and a small (< 2 mm) device architecture. Our sharpening method exhibits the carbon fibers of smallest known length to be sharpened, making carbon fibers suitable for chronic interfacing with small diameter autonomic nerves. We also show that long, sharpened, individuated carbon fibers are capable of self-insertion to depths of the brain previously only reached with insertion assistance. Our *in vivo* tests of SCFs show insertion into neural structures of varying stiffness and depths. Our benchtop experiments test the durability of carbon fibers embedded in silicone and the capability to survive many repeated cycles of bending. This suggests that subcellular carbon fiber electrodes with a stress-relieving backplane are a suitable electrode material for chronic residence in peripheral structures. We test the unique components of the array in the rat CVN, feline dorsal root ganglia (DRG), and rat motor cortex. We demonstrate that SCF arrays record spontaneous and physiologically-relevant neural clusters. Additionally, we verify that the small device architecture enables surgical handling while accessing neural structures of differing geometries. This work showcases a versatile, unique set of capabilities for carbon fiber arrays in neural interfacing and indicates the potential for chronic use.

## II. Methods

### A. Fabrication of Carbon Fiber Flex Array

Acute neural recordings from the rat CVN and feline DRG were collected using carbon fiber electrode arrays previously detailed in Patel et al., 2020 [38]. In this application, the carbon fibers were cut to 150–200 μm prior to sharpening.

### B. Fabrication of Carbon Fiber Silicone Array (CFSA)

This study exhibits a novel carbon fiber silicone array (CFSA) featuring carbon fibers partially embedded in silicone (Fig. 1). The overall array size is 1.75 mm x 500 μm x 750 μm (length x width x thickness). The array is built on a custom polyimide PCB (MicroConnex, Snoqualmie, WA, USA), 450 μm x 6.81 mm x 50 μm in size. A 1.5 mm long exposed section of the PCB contains 4 pairs of 430 μm-long gold traces with 65 μm pitch that are positioned 310 μm apart (Fig. 1B). Each trace pair is electrically connected to a 90 μm-diameter gold-plated via. During fabrication, the polyimide PCB is slotted into a custom aluminum holder (Fig. 1A) (Protolabs, Maple Plain, MN, USA) and secured with double-sided carbon tape.

**Fig. 1.**
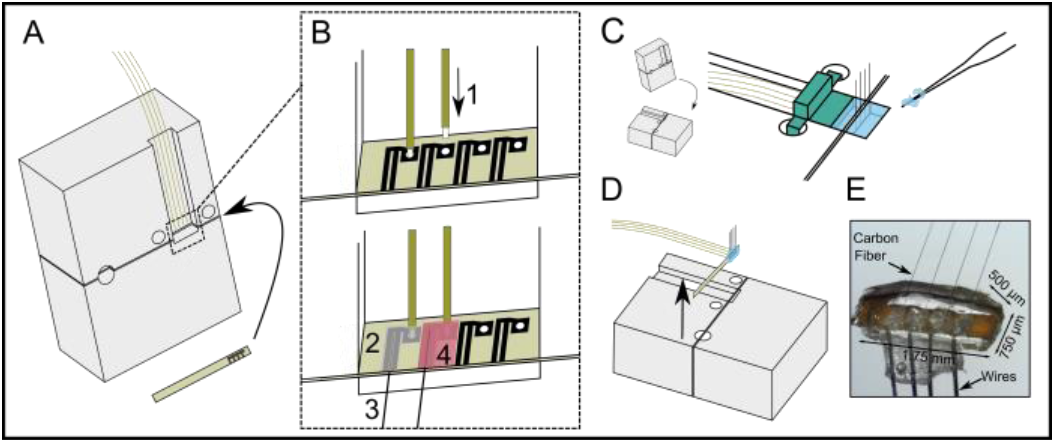
Fabrication steps of carbon fiber silicone array (CFSA). A) A custom polyimide PCB (gold) is slotted into a custom aluminum holder (grey) and four polyimide-insulated wires (gold) are positioned above the area of interest on the polyimide PCB. B) The wires and carbon fibers are electrically connected and secured to the polyimide PCB. B1) The tip of each wire is exposed from polyimide insulation and placed into the gold-plated vias in the polyimide PCB. B2) Silver epoxy is applied to the pair of connected gold traces and the adjoining gold-plated via. B3) Bare carbon fibers are placed in the silver epoxy between the pair of gold traces, and then cured. B4) Insulating epoxy is applied over the electrical connections, and then cured. C) The aluminum holder is laid flat and a 3-D printed device (green) is placed over the wires to create a well for the silicone. Silicone (blue) is applied with pulled glass capillaries and then cured. D) The polyimide PCB with attached wires, carbon fibers, and silicone is lifted from the holder. E) A picture of the CFSA with the excess tails of polyimide PCB on either side of the silicone removed.

A custom interface PCB was designed to interface with either a 32 or 16-channel Omnetics connector (A79024-001 or A79040-001, Omnetics, Minneapolis, MN, USA) and four 10 cm-long wires (50.8 μm diameter 35N LT, coated with polyimide to 60.9 μm, Fort Wayne Metals, Fort Wayne, IN, USA). The Omnetics connector and four wires were soldered to the interface PCB and secured with 2-part quick curing epoxy. The other wire end, with 75 μm uninsulated, was slotted into a gold-plated via in the polyimide PCB (Fig. 1B-1). Silver epoxy was applied to the junction of the PCB and wire and the adjoining pair of gold traces (Fig. 1B-2). Individual carbon fibers were placed in the silver epoxy between the paired gold traces (Fig. 1B-3). After the silver epoxy was cured, a layer of insulating epoxy (353ND-T, Epoxy Technology, Inc., Billerica, MA, USA) was applied over traces and wire junctions and cured (Fig. 1B-4).

A custom 3D-printed piece was attached to the aluminum holder to create a well around the CFSA for the silicone molding (Fig. 1C). Pulled glass capillaries shuttled the degassed medical grade silicone (A-103, Factor II, Lakeside, AZ, USA) into the 200 μm-wide areas on either side of the polyimide PCB and across the sides between the fibers. The silicone was cured at 110° C for 20 minutes and the 3D printed wall was removed. The device was coated with approximately 800 nm of parylene-c. The polyimide PCB, with attached silicone, fibers, and wires, was removed from the aluminum holder by gently pulling on the free end of the PCB with forceps (Fig. 1D). Once removed, the carbon fibers were trimmed to length (500–1100 μm based on application) with microsurgical scissors prior to electrode tip processing (Fig. 1E).

### C. Sharpening of Carbon Fiber Electrode Tip

We adapted a heat-based sharpening method using a butane torch, seen in previous demonstrations of carbon fibers in long fiber bundles [34] or relatively long individuated fibers [37], to work with short individuated fibers (200–250 μm) suitable for dwelling within 300-500 μm-diameter nerves. Our method uses the reflective properties of water to precisely align short fibers with the water’s surface for sharpening while the body of the array is kept safely under water (Fig. 2).

**Fig. 2.**
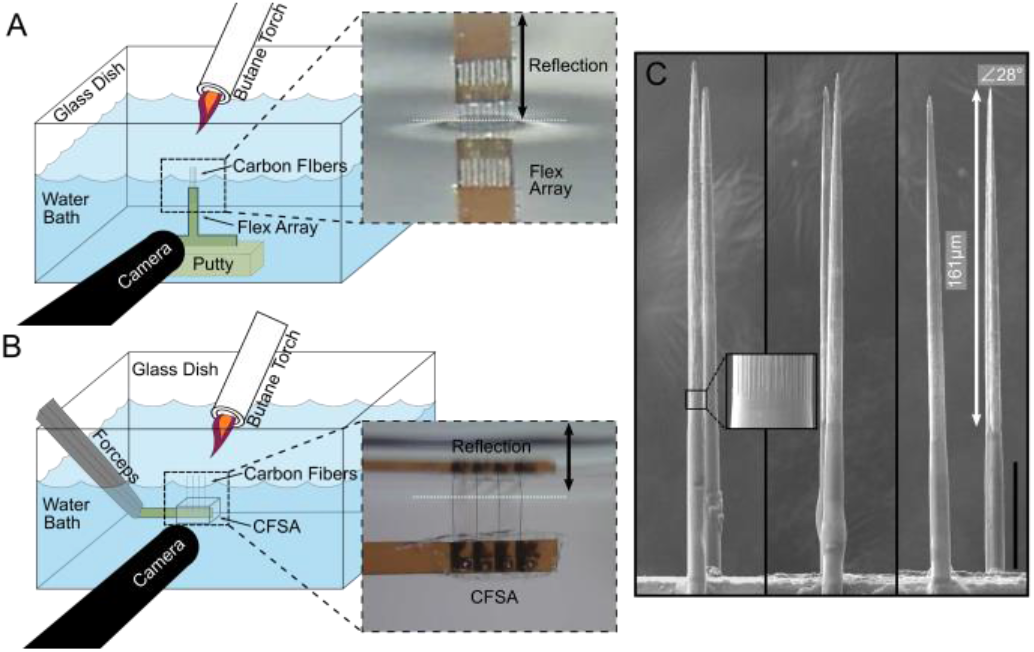
Blowtorch schematic and sharpened fibers. A) Flex Arrays are secured with putty to the water dish base. The water level is adjusted until the carbon fiber tips touch their reflection on the underside of the water surface (inset, black arrow denotes reflection). B) CFSAs are held with forceps and lowered into the water dish until the carbon fiber tips touch their reflection on the underside of the water surface (inset, black arrow denotes reflection). With both arrays, a pen camera located outside the water dish views the carbon fiber reflection under the water surface to confirm alignment at the water surface. A butane flame is passed over the water surface to sharpen the carbon fibers. C) Sharpened fibers have a smooth transition from parylene-c to bare carbon (inset). The representative SCF on the right has 161 μm of carbon exposed from parylene-c (white arrow) and a tip angle of 28 degrees. Scale bar is 50 μm.

Flex Arrays and CFSAs underwent a similar sharpening procedure using different holding mechanisms. The Flex Array was secured with putty to the base of a glass dish (Fig. 2A). The water level was adjusted with a pipette until the array was completely submerged. The small, lightweight nature of the CFSAs required it be held by nerve forceps (ASSI.NHF0.5, Accurate Surgical & Scientific Instruments Corp., Westbury, NY, USA) duing sharpening (Fig. 2B). The tail of the CFSA was held at a 45 degree angle by forceps, which were secured to a stereotactic frame. The surface of the CFSA was parallel to the water surface with carbon fibers pointing up as it was lowered into the water until submerged.

An endoscopic camera (MS100, Teslong, Shenzhen, China) aligned outside the glass dish was used to view the carbon fibers. The camera was tilted up beneath the water level to visualize the reflection of the fibers on the underside of the water surface. The water level was adjusted until fibers appeared to touch the reflection on the underside of the water surface (Fig. 2A, B insets), at which point a microtorch (3 mm diameter flame, MT-51B, Master Appliance, Racine, WI, USA) was passed over the fibers for 10–20 s (Fig. 2C). Sharpened carbon fibers (SCFs) were confirmed by visual inspection. Multiple passes of the microtorch were occasionally necessary.

### D. Scanning Electron Microscopy (SEM) Imaging

Scanning Electron Microscopy (SEM) images were used to characterize SCFs. Images were collected in either a Nova Nanolab 200 DualBeam SEM (FEI, Hillsboro, OR, USA) or a MIRA3 SEM (Tescan Orsay Holding, Brno-Kohoutovice, Czech Republic). An accelerating voltage of 2 kV or 3 kV and a current of 0.21 nA or 24 pA was used on the Nova or MIRA3, respectively. Both SEMs used an Everhart-Thornley detector for high-vacuum secondary electron imaging. Arrays were mounted on standard SEM pin stub mounts using carbon tape and gold sputtered for 60–120.

### E. Electrochemical Deposition of Polymer Coating

To lower impedance, a solution of 0.01 M 3,4-ethylenedioxythiophene (483028, Sigma-Aldrich, St. Louis, MO, USA) and 0.1 M sodium p-toluenesulfonate (152536, Sigma-Aldrich, St. Louis, MO, USA) was electrodeposited onto the exposed carbon of the SCFs by applying 600 pA/channel for 600 s to form poly(3,4-ethylene-dioxythiophene):sodium p-toluenesulfonate (PEDOT:pTS) [30], [35].

### F. Electrochemical Impedance Spectroscopy (EIS) and Cyclic Voltammetry (CV) Analysis

EIS and CV measurements were collected to characterize electrode fabrication steps and post-surgery viability. Three-electrode measurements were performed with a PGSTAT12 Autolab potentiostat (Metrohm / Eco Chemie, Utrecht, Netherlands) and vendor-supplied Nova software using the same setup and analysis methods described in Welle et al., 2020 [36]. Fibers were rinsed in deionized water after measurement.

### G. Bend Testing of Silicone-Embedded Carbon Fibers

Carbon fibers were embedded in medical grade silicone to test their bending characteristics. Silicone was held in oval depressions milled into an aluminum baseplate (Protolabs, Maple Plain, MN, USA) and degassed in a vacuum chamber for 20 minutes at −0.09 MPa. Eight bare carbon fibers at a 152.4 μm pitch were aligned and submerged into silicone by 300–400 μm. After curing on a hotplate for 20 minutes at 110° C, the silicone-CF testing devices were gently removed with forceps. The silicone surface of some devices was slightly curved, but this was deemed insignificant for our purposes. In a small number of experiments, CFSAs were tested to verify the amount of silicone needed above the edge of the polyimide PCB to exhibit the same properties as CFs embedded purely in silicone [39].

A 0.75 mm diameter glass capillary a linear actuator (M-235.5DD, PI, Auburn, MA, USA) was repeatedly passed over the carbon fibers for bend testing. The glass capillary was 40–60 μm above the silicone surface and moved at a velocity of 17.09 mm/s and acceleration of 4.88 mm/s^2^. One pass was the movement of the glass capillary from one end of the line of carbon fibers to another in a single direction. The percent of broken fibers on each device was collected at some combination of the experimental increments of 200, 1000, 2000, 6000, 8000, 10000, 20000, 30000, 40000, and 50000 passes.

### H. Animal Surgery

All animal procedures were approved by the University of Michigan Institutional Animal Care and Use Committee (PRO00009525). Electrophysiological recordings were collected from three locations in non-survival surgical procedures: rat CVN, feline DRG, and rat brain motor cortex. Prior to all surgeries, ground and reference wires (AGT05100, World Precision Instruments, Sarasota, FL, USA) were soldered to the interface PCB and exposed from insulation by approximately 1 cm on the other end. The polyimide PCB tail was clamped by nerve forceps such that the fibers pointed away from the forceps toward down the cavity.

### I. Acute Rat CVN Surgery

The non-survival CVN procedures were performed on male Sprague-Dawley rats (0.36–0.62 kg, Charles River Laboratories, Wilmington, MA, USA), as detailed in Jiman et al., 2020 (N=22 for companion study, +1 additional animal in current study), using Flex Arrays (150–250 μm length SCFs). Isoflurane (Fluriso, VetOne, Boise, ID, USA) was used for anesthetic induction (5%) and maintenance (2–3%). Rats were placed on a heating pad and vitals were monitored. The CVN was accessed through a midline ventral cervical incision. Approximately 10 mm of CVN was isolated from the carotid artery and surrounding tissue. The CVN was lifted (~2 mm) on to a custom 3D-printed nerve-holder. The Flex Array was positioned over the elevated CVN using a 3-axis micromanipulator (KITE-R, World Precision Instruments, Sarasota, FL, USA) secured to an optical breadboard. The ground wire for the Flex Array was inserted subcutaneously in the cervical area and the reference wire was placed in the tissue of the cervical cavity. An endoscopic camera was aligned at the edge of the nerve-holder to visualize the carbon fibers during insertion. The heating pad and dissection microscopy were disconnected to reduce electrical noise. The nerve was rinsed with saline (0.9% NaCl, Baxter International, Inc., Deerfield, IL, USA) and the Flex Array carbon fibers were lowered and inserted into the CVN.

### J. Acute Feline DRG Surgery

Three adult, domestic, short-hair male felines (4.0–5.6 kg, 1.3-1.4 years, Marshall BioResources, North Rose, NY, USA) were used for the DRG experiments with Flex Arrays (150–250 μm length SCFs). One feline (5.2 kg, 1.2 years) was used for the DRG experiment with a CFSA (375–475 μm length SCFs). Flex Arrays were previously used in CVN experiments. The animals were anesthetized with an intramuscular injection of ketamine (6.6 mg/kg), butorphanol (0.66 mg/kg), and dexmedetomidine (0.011 mg/kg). Animals were intubated and maintained on isoflurane (0.5–4%) and vitals were monitored (respiratory rate, heart rate, end-tidal CO_2_, O_2_, perfusion, temperature, and intra-arterial blood pressure). For fluid infusion and pressure monitoring, a 3.5 Fr dual-lumen catheter was inserted to the bladder through the urethra. A midline dorsal incision was made to expose the L7 to S3 vertebrae. A laminectomy was performed to access the S1–S2 DRG. After experimentation, animals were transitioned from isoflurane to intravenous alpha-chloralose (70 mg/kg induction and 20 mg/kg maintenance) for the remainder of the experiment. Buprenorphine was provided subcutaneously every 8 to 12 hours as analgesia.

The Flex Array or CFSA was held in place with a clamp or nerve forceps and manually lowered into the surgical site using a micromanipulator. The ground wire was placed subcutaneously at a distant location and the reference wire was placed near the spinal nerve. Insertion into the S1 or S2 DRG was visualized with an endoscopic camera. After insertion, impedances were recorded at 1 kHz. Brushing trails with the Flex Arrays consisted of brushing of the scrotum with a cotton-tip applicator for 10 seconds following a period of no brushing for 10 seconds, for 60 seconds total. For bladder trials with the Flex Array, saline was infused into an empty bladder at 2 mL/min until either an elevated bladder pressure occurred or urine leaking was observed. CFSA trials consisted of exploratory perineal brushing and baseline recordings.

### K. Acute Rat Cortical Surgery

Four adult male Sprague Dawley rats (0.3–0.4 kg, Charles Rivers Laboratories, Wilmington, MA, USA) were used for the acute cortical experiments with CFSAs (75–1075 μm length SCFs). Two animals were excluded due to surgical complications unrelated to device features. Animals received carprofen (5 mg/kg) and were anesthetized with ketamine/xylazine (90/10 mg/kg) and maintained with ketamine (30 mg/kg). Rat vitals were monitored during surgery. Surgical preparation is detailed in Welle et al., 2020. A 2.5 mm by 2.5 mm craniotomy was drilled over the right hemisphere motor cortex. The CFSA was attached to a stereotactic manipulator and aligned over the craniotomy along the anterior-posterior axis. The ground and reference wires were wound around the grounding bone screw. A dural slit was made in the center of the craniotomy. The CFSA was manually lowered and the fibers were driven into the brain until the final target depth (0.775–1.180 mm) in layer V [41] was reached.

### L. Tip Electrophysiology Recording

Neural recordings from the rat CVN and feline DRG were collected using a Grapevine Neural Interface Processor (Ripple LLC, Salt Lake City, UT, USA). Electrophysiological signals were recorded at a sampling rate of 30 kHz. Impedances were measured during rat CVN experiments at 1 kHz in saline before the procedure and in the nerve immediately after insertion. Impedances were measured during the majority of feline DRG experiments at 1 kHz immediately after insertion. At least 5 minutes, 10 minutes, or 1 minute of recordings were obtained for the rat CVN, feline DRG bladder experiments, and feline DRG brushing experiments.

Brain electrophysiology recordings were collected using a ZC16 headstage, RA16PA pre-amplifier, and RX5 Pentusa base station (Tucker-Davis Technologies, Alachua, FL, USA). Data was sampled at a rate of 25 kHz, high-pass filtered at 2.2 Hz, and anti-alias filtered at 7.5 kHz. Each recording session lasted at least 3 minutes.

### M. Analysis of Neural Recordings

Principle component analysis of neural recordings was conducted in Offline Sorter (OFS, Plexon, Dallas, TX, USA) to isolate neural clusters. The electrophysiology signals were filtered with a band-pass filter at 300–10,000 Hz and manually thresholded below the noise floor. A trained operator manually identified neural clusters from the CVN and DRG recordings.

Clusters from the brain were identified in OFS using the semi-automated method described in Welle et al., 2020. Sorted clusters were identified by unique amplitude, waveform shape, inter-spike interval, and response to physiological signals, such as breathing rate. MATLAB was used to analyze the sorted clusters. Firing rates of bladder clusters were calculated with a bin duration of 1 second and correlated to bladder pressure until the maximum bladder pressure [42]. The signal-to-noise ratio (SNR) was calculated using the mean peak-to-peak amplitude (V_pp_) of a sorted cluster and the standard deviation of at least 500 ms of noise from the respective recording [SNR = V_pp_ / (2 * standard deviation of noise)]. When appropriate, values are presented as mean ± standard deviation (SD).

## III. Results

### A. Analysis of Sharpened Carbon Fibers

A blowtorch method for sharpening <500 μm linear carbon fiber arrays was developed to facilitate consistent insertion success in peripheral targets (Fig. 2A, B). Example SCFs attached to Flex Arrays are shown in Fig. 2C. Prior to blowtorching, carbon fibers were manually cut using microsurgical scissors to roughly 200 μm in length. After blowtorching, SEM analysis confirmed the average length of SCFs as 223.7 ± 18.9 μm (N=32 fibers). SCFs exhibited an average tip angle of 72.3 ± 33.5 degrees with a 146.8 ± 17.7 μm length of carbon exposed from parylene-c insulation, constituting an active recording site of 2734.5 ± 402.5 μm^2^ surface area (Fig. 2C). The parylene-c transition between the exposed and insulated carbon appeared smooth on all imaged fibers (Fig. 2C inset).

After sharpening, the electrodes were coated PEDOT:pTS [35]. The SCFs coated with PEDOT:pTS showed a large shift in impedance magnitude at lower frequencies (Fig. 3A). SCFs of 200 μm length without PEDOT:pTS coating exhibited an average 1 kHz impedance (|Z|_1kHz_) of 334.8 ± 243.2 kΩ and a median |Z|_1kHz_ of 298.6 kΩ (N=574 fibers). Once coated with PEDOT:pTS, SCFs exhibited an average |Z|_1kHz_ of 32.3 ± 55.7 kΩ and a median |Z|_1kHz_ of 12.4 kΩ (N=565 fibers) (Fig. 3B). Therefore, coating the active SCF recording site with PEDOT:pTS contributed to a 90% decrease in average |Z|_1kHz_ (Fig. 3B). Correspondingly, the cathodal charge storage capacity of SCFs increased from 15.2 ± 5.4 μC/cm^2^ to 255.0 ± 148.6 μC/cm^2^ (N=64 fibers), calculated at a sweep rate of 1 V/s, after PEDOT:pTS deposition (Fig. 3C).

**Fig. 3.**
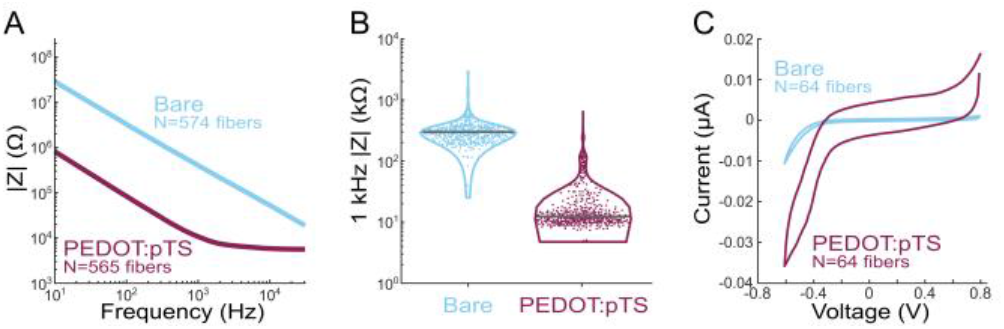
Electrochemical impedance spectroscopy and cyclic voltammetry (CV) analysis of SCFs. A) Impedance magnitude frequency spectrogram of bare SCFs (blue, N=574) and PEDOT:pTS coated SCFs (magenta, N=565). B) Violin plot of 1 kHz impedance magnitudes of bare (N=574) and PEDOT:pTS coated (N=565) SCFs. Mean 1 kHz impedance magnitude was 334.8 ± 243.2 kΩ and 32.3 ± 55.7 kΩ for bare and PEDOT:pTS coated SCFs. C) CV curves of bare (N=64) and PEDOT:pTS coated (N=64) SCFs. The cathodal charge storage capacity increased from 15.2 ± 5.4 μC/cm2 to 255.0 ± 148.6 μC/cm2 once SCFs were coated with PEDOT:pTS.

### B. Durability Testing of Silicone-Embedded Carbon Fibers

We harnessed the intrinsic compliance of carbon fibers through partial embedding in medical grade silicone [30], [39] to relieve strain at the SCF-polyimide PCB junction, therefore minimizing the eventual breakage seen on Flex Arrays after repeated insertions. Bend tests were conducted on silicone-embedded carbon fibers, with and without attachment to an embedded polyimide PCB, using a 0.75 mm diameter glass capillary (Fig. 4A, C, Supplemental Video 1, Supplemental Video 2). On average, carbon fibers deflected 50.9 ± 6.2 degrees from the vertical position (N=23 fibers, 3 devices), representatively shown in Fig. 4B. The maximum measured fiber deflection from vertical without fracture was 71.6 degrees. We observed that an average of 93.8% fibers (N=58/62) did not break after 2,000 bends (passes of the capillary) during continuous fatigue testing (Fig. 4D). After 50,000 passes, 87.5% of fibers remained unbroken (N=14/16). This durability was similarly observed on carbon fibers embedded in at least 150 μm of silicone when attached to an embedded polyimide PCB (Woods et al., 2020, Supplemental Video 2).

**Fig. 4.**
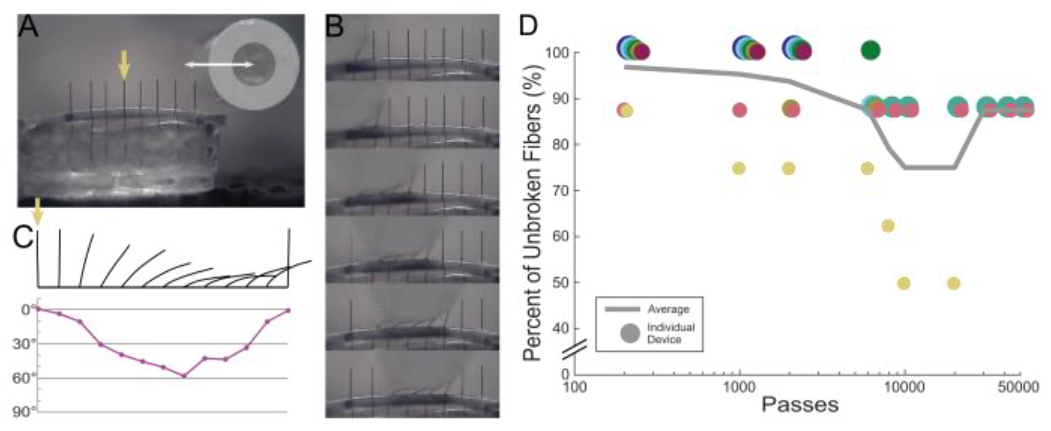
Bend test of silicone-embedded carbon fibers. A) Bend test setup with a silicone carbon fiber testing device and a 0.75 mm diameter glass capillary (outlined in light grey). The glass capillary passes over the carbon fibers. A complete run from end to end counts as one pass. B) Still images of bent carbon fibers from a video shown in (A) and in Supplemental Video 1 during one pass. C) Top: Outlines of a single fiber, indicated in (A) by the yellow arrow, bending under the glass capillary. Bottom: Corresponding angle measurements from the outline of the fiber in each still image. On average, carbon fibers deflected 50.9 ± 6.2 degrees from the vertical position (N=23 fibers, 3 devices). D) Bend test data showing the percent of unbroken fibers plotted against the number of glass capillary passes. Each device contains 8 fibers and is shown as an individually colored circle. The average across all devices is shown as the solid line. After 50,000 passes, 87.5% of fibers remained unbroken.

### C. Acute Electrophysiology Recordings from Rat CVN

We tested the ability of SCFs to insert into and record from small diameter (300–500 μm) rat CVN. Six Flex Arrays with SCFs of 200–250 μm length, as represented in Fig. 1C, were inserted into the left CVN of 22 Sprague-Dawley rats (Fig. 5E, Supplemental Video 3). Detailed characterization of the population recordings is presented in a companion study by Jiman *et al.* 2020 [40]. In all experiments, we visually observed successful insertion of all SCFs into the CVN after an average of 2.3 insertion attempts. The functional SCFs (defined as |Z|_1kHz_ < 1 MΩ) had an average |Z?1kHz of 70.8 ± 81.9 kΩ once inserted into the CVN (N=326 fibers). Neural activity was recorded on 51.2% of the functional SCFs. Manual sorting and a mixture of Gaussians sorting methods were applied in principal component space to classify neural activity into clusters of distinct bipolar waveform shapes. Across the 326 functional SCFs, 174 sorted neural clusters were detected. The neural clusters had a mean V_pp_ of 30.7 ± 11.43 μV and a mean SNR of 3.52 ± 1.0. We observed that 19.8% of functional SCFs recorded neural clusters with SNRs > 4. The maximum mean V_pp_ was found to be 91.72 μV.

**Fig. 5.**
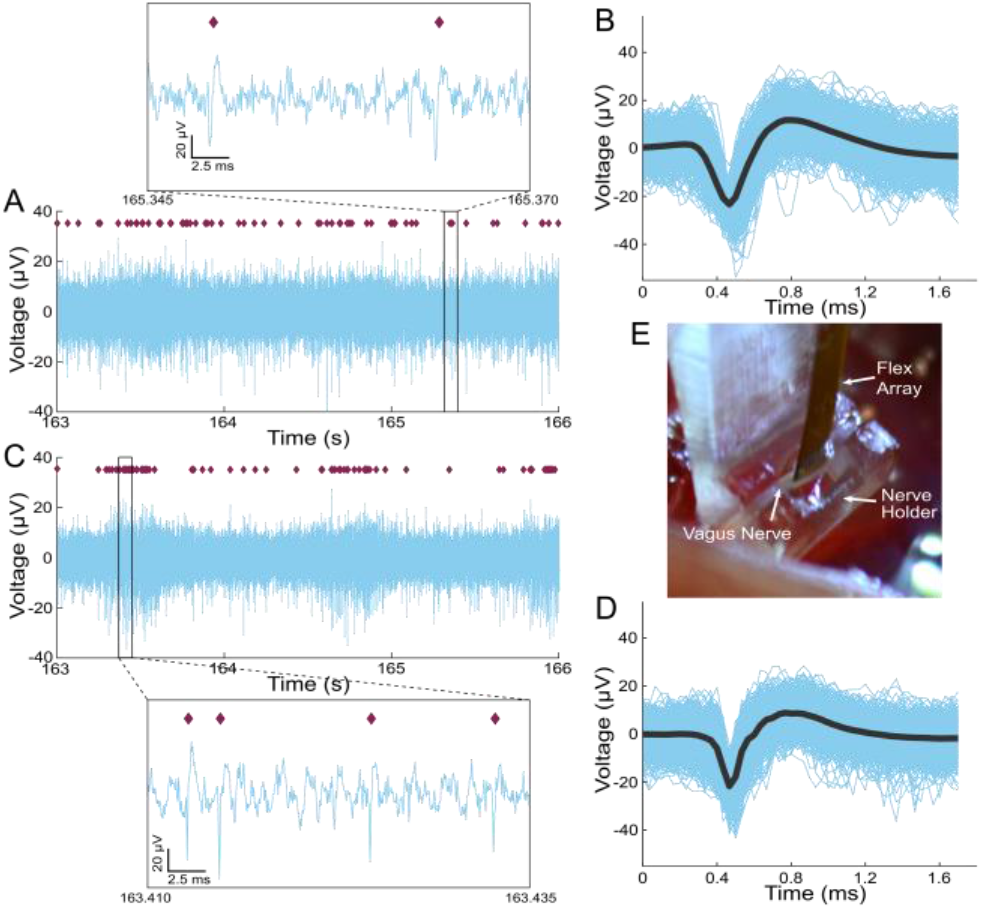
Neural activity recorded from a rat CVN with SCFs on a Flex Array, representative of the larger data set shown in Jiman et al., 2020. A and C) Three seconds of filtered data recorded on two SCFs. An inset of each raw trace shows 25 μs of data. B and D) Corresponding neural cluster indicated in (A) and (C) by purple diamond with the mean cluster waveform shown by grey solid line. B) The mean V_pp_=139.9 ± 26.4 μV. D) The mean V_pp_=122.8 ± 24.3 μV. E) Surgical setup showing the CVN lifted by the nerve holder with Flex Array aligned above it.

Representative electrophysiological recordings of spontaneous CVN activity is shown in Fig. 5. Sixteen SCFs were determined functional and had an average |Z|_1kHz_ of 25.3 ± 10.9 kΩ after insertion into the CVN. Sorting analysis identified neural clusters on 3 of 16 functional SCFs in this example experiment. Neural data from 2 functional SCFs are shown in Fig. 5A and C. The corresponding neural cluster from each functional SCF had a mean V_pp_ of 139.9 ± 26.4 μV and 122.8 ± 24.3 μV (Fig. 5B, D). Across all 3 SCFs, we found that the waveform V_pp_ of clusters ranged from 18.5 μV to 108.5 μV. The sorted neural clusters had a mean V_pp_ of 41.8 ± 6.8 μV and an average median V_pp_ of 41.0 μV. Clusters from this representative experiment had a mean SNR of 4.1 ± 0.08.

### D. Acute Electrophysiology Recordings from Feline DRG

We sought to discover whether it is possible to insert SCFs of short length (150–250 μm) into feline DRG with minimal external force during insertion. We inserted 16-channel Flex Arrays with 150–250 μm length SCFs into the S1 DRG of 3 felines (Fig. 7A, B) and a 4-channel CFSA with 375–475 μm length SCFs into the S2 DRG of 1 feline (Fig. 6A, B). In each experiment, the array was manually lowered with a micromanipulator until SCFs were visually inserted in tissue. Functional SCFs (|Z|_1kHz_ < 1 MΩ) had an average |Z|_1kHz_ of 26.9 ± 7.4 kΩ in the DRG immediately after insertion (N=30 fibers).

**Fig. 6.**
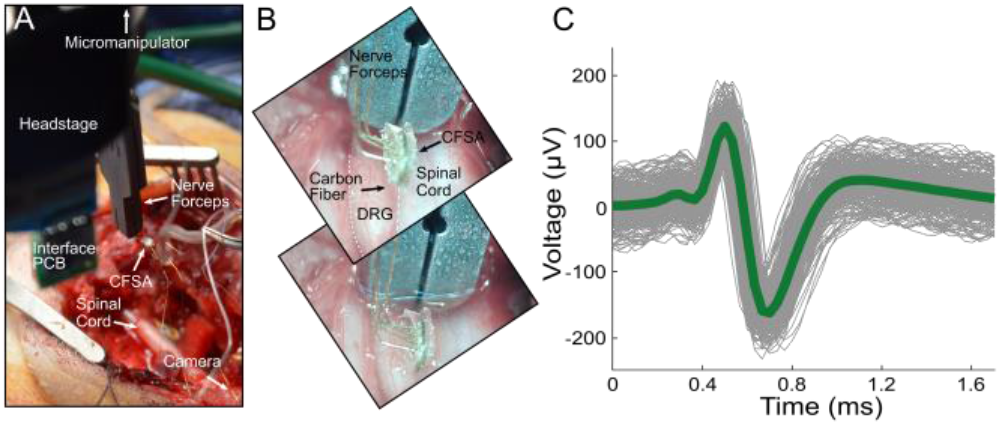
CFSA experiment in feline DRG. A) Surgical setup showing the CFSA, held by nerve forceps, and interface PCB connected to a headstage lowered to the DRG with micromanipulator. The DRG is located below the visible spinal cord. The camera is positioned parallel to the spinal cord to the right of the surgical cavity. B) CFSA, held by nerve forceps, as seen by the camera before (top) and after (bottom) insertion into the DRG (white dashed line). C) Spontaneous neural cluster and mean waveform shape (green line) recorded in the DRG with the CFSA (mean V_pp_=298.8 ± 15.5 μV, SNR=6.5).

**Fig. 7.**
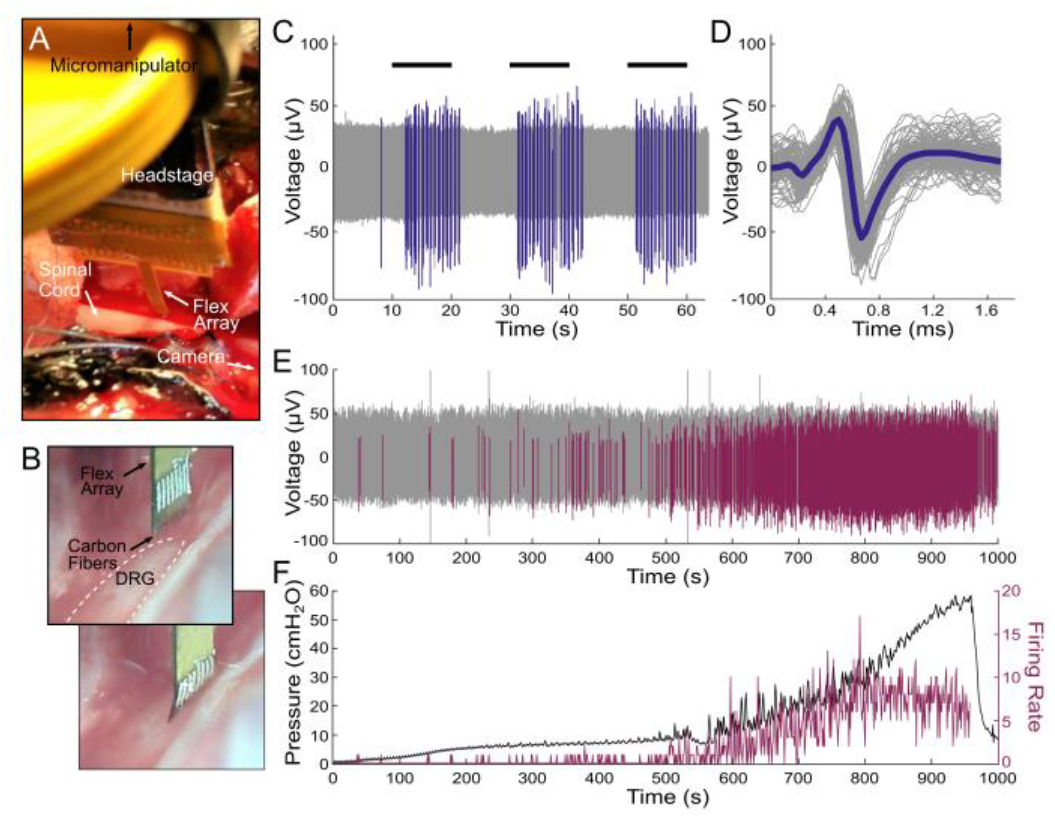
Flex Array experiments in feline DRG. A) Surgical setup showing the Flex Array connected to a headstage lowered to the DRG with a micromanipulator. The DRG is located above the visible spinal cord. The camera is positioned parallel to the spinal cord to the right of the Flex Array. B) Flex Array as seen by the camera before (top) and after (bottom) insertion into the DRG (white dashed lined). C) Neural waveforms in response to scrotal brushing (horizontal black lines) are shown in blue. D) The brushing cluster from (C) with mean waveform shown by thick blue line (mean V_pp_=108.7 ± 11.0 μV, SNR=6.0). E) Neural waveforms associated with bladder filling are colored purple. F) Bladder pressure (left axis) and firing rate (right axis) of bladder cluster in (E), with correlation coefficient of 0.83.

Despite the large recording surface area, neural clusters were observed on 32 of 52 total inserted SCFs (N=4 felines; 3 Flex Arrays; 1 CFSA), and 27 of 30 functional (|Z|_1kHz_ < 100 kΩ) SCFs (N=3 felines; 2 Flex Arrays; 1 CFSA). Impedance data was not recorded on one feline and excluded from the functional SCF count. Our offline sorting analysis of recorded neural activity identified 73 neural clusters across all feline experiments with an average of 1.8 ± 0.9 clusters per SCF. Clusters were classified as either spontaneous or driven. Spontaneous neural activity was sorted into clusters associated with breathing (breathing clusters), clusters unlinked to obvious physiological signal (spontaneous clusters, Fig. 6C), and clusters containing multiple units (multi-unit clusters). Driven clusters responded to the experimental variables of perineal or scrotal brushing (brushing clusters, Fig. 7C, D) or saline-infusion of the bladder (bladder clusters, Fig. 7E, F). Across all experiments, we classified 16.2% of clusters as breathing, 43.2% as spontaneous, and 16.2% as multi-unit, 14.9% as brushing, and 9.5% as bladder clusters.

Breathing clusters depicted rising and falling amplitudes, in addition to bursting patterns. We believe the observed changes in amplitude were likely due to micromotion shifts in SCF position with respect to the neuron as the animal’s body shifted slightly with each breath. The V_pp_ of breathing clusters ranged from 41.4 μV to 186.7 μV, with a mean V_pp_ of 84.4 ± 16.8 μV and a mean SNR of 7.0 ± 3.7 (N=12 clusters; 3 Flex Arrays). Spontaneous clusters did not obviously track with breathing or other physiological signals. The mean V_pp_ of spontaneous clusters was 115.9 ± 16.6 μV and a mean SNR of 7.9 ± 6.0 (N=32 clusters; 3 Flex Arrays; 1 CFSA). Fig. 6C shows an example spontaneous cluster recorded from the CFSA with a mean V_pp_ of 298.8 ± 15.5 μV and SNR of 6.5. Multi-unit clusters contained clear neural activity, but we were unable to differentiate spontaneous clusters. Multi-unit clusters had a mean V_pp_ of 34.2 ± 6.3 μV and a mean SNR of 4.7 ± 1.2 (N=12 clusters; 1 Flex Array).

To further verify that spikes were neural in origin, we recorded neural activity in response to cutaneous brushing and bladder filling. In 2 of 3 experiments with Flex Arrays recording from the S1 DRG, we recorded neural activity corresponding to the cutaneous response to scrotal brushing. We observed 10 clusters on 8 SCFs with bursting activity during the 10 second brushing intervals (Fig. 7C). The corresponding brushing cluster, shown in Fig. 7D, has a mean V_pp_ of 108.7 ± 11.0 μV and SNR of 6.0. The mean V_pp_ of all brushing clusters was 152.4 ± 13.5 μV and the mean SNR was 7.1 ± 4.5 (N=10 clusters; 2 Flex Arrays). We also recorded a single brushing cluster in response to perineal brushing on a CFSA in one experiment. The mean V_pp_ of the cluster was 219.5 ± 54.1 μV and the mean SNR was 18.9 (N=1 cluster; 1 CFSA)

We also observed neural activity that contained physiologically-relevant signals of bladder pressure in response to bladder filling. Across all experiments, the firing rates of 7 neural clusters were found to correlate with bladder pressure during the bladder filling procedure. A representative bladder cluster is shown in Fig. 7E and 7F. Correlation coefficients between bladder pressure and firing rate were calculated for each bladder cluster. The correlation coefficients were between 0.23 and 0.82, with a median of 0.70. The mean correlation coefficient of the 7 clusters with their respective bladder pressure was 0.61 ± 0.21. The V_pp_ of the bladder clusters were between 32.1 μV and 190.4 μV, with a mean V_pp_ of 86.8 ± 14.1 μV and a mean SNR of 10.3 ± 6.3 (N=7 clusters, 3 Flex Arrays).

We observed that several clusters recorded on the Flex Array appeared on multiple adjacent SCFs. These units fulfilled our metrics for being a sortable cluster in terms of amplitude, signal width, and inter-spike-interval, but appeared concurrently on multiple consecutive channels. In one instance, a breathing cluster was seen on four SCFs in a 2 x 2 geometry on a Flex Array, spanning a distance of 141 μm between diagonal SCFs.

### E. Acute Electrophysiology Recordings from Rat Cortex

SCFs on CFSAs of length 750 μm to 1075 μm were found to self-insert into dura-free brain tissue from depths of 775 μm to 1180 μm (Fig. 8B, Supplemental Video 4). In total, 17 sortable single units were recorded from 6 SCFs across 3 devices during 7 independent insertions into the brain (Fig. 8A). The mean V_pp_ was found to decrease as cortical depth increased (Fig. 8 table). The overall mean V_pp_ was 80.7 ± 27.5 μV and the largest recorded unit amplitude was 126 μV.

**Fig. 8.**
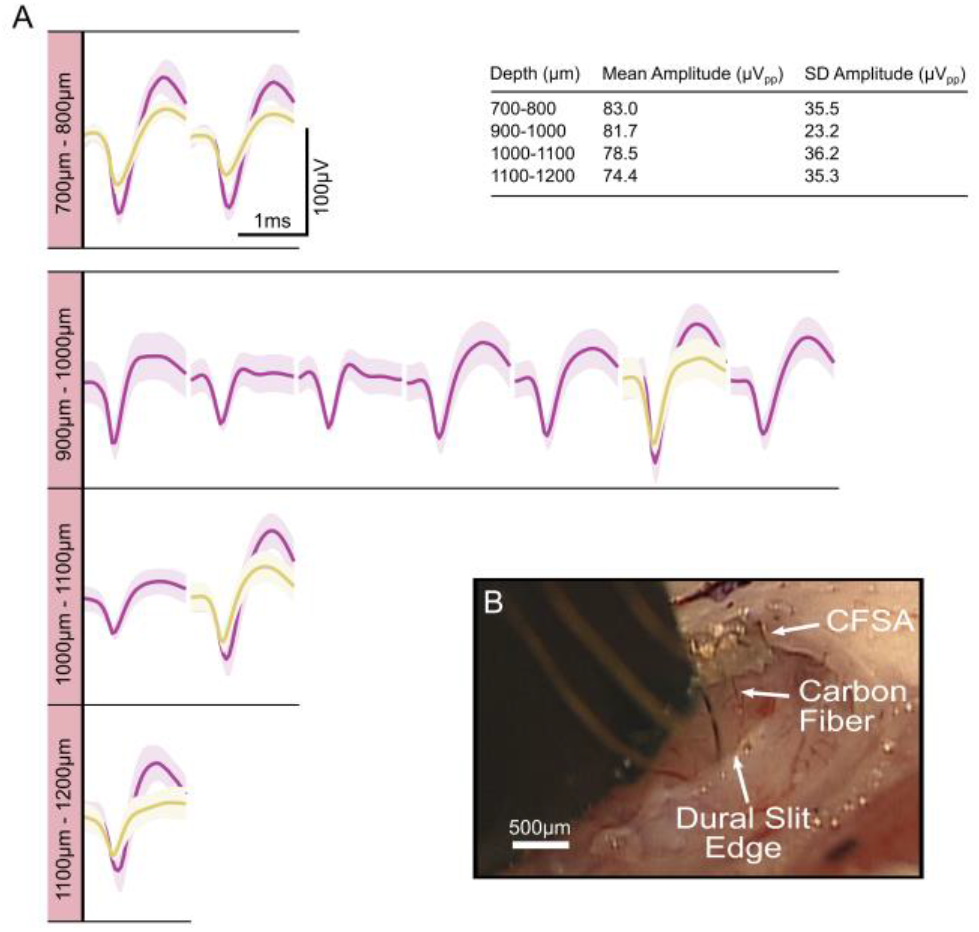
Electrophysiology recorded from SCFs on CFSAs in the rat motor cortex. A) Neural units recorded at varying depth increments. The units were recorded on 6 SCFs across 3 devices during 7 independent insertions into the brain. Each unit, or overlapping units, was recorded from a single SCF during one insertion. The inset table shows a summary of the mean amplitude and standard deviation of neural units at each depth. B) Surgical setup of CFSA insertion into the brain through the dural slit.

## IV. Discussion

Here, we demonstrate the capabilities of SCF arrays for interfacing with the central and peripheral nervous systems. The work presented here represents, to our knowledge, the first stiff, penetrating, sub-cellular electrode array with a durable, stress-relieving backplane.

SCFs easily penetrate into multiple neural structures of varying stiffness and geometry. Initial insertion attempts in the small diameter (300–500 μm) rat CVN with conventional blunt-tipped carbon fiber electrodes [36] yielded unsuccessful insertion, and surgeons often resorted to slitting the epineurium. However, short (< 250 μm) SCFs consistently inserted into the CVN. Successful insertion was also achieved in the feline DRG, which has a thick epineurium layer of 50–100 μm [43], [44], without the assistance of the pneumatic inserter [15]. In the brain, blunt-tipped carbon fibers previously relied upon insertion assistors, such as temporary stiffening agents or a silicon backbone, to insert into dura-free brain at lengths greater than 500 μm [35], [36]. Despite a 6.8 μm diameter, SCFs inserted without assistance into brain at lengths of 1075 μm.

Silicone-embedded carbon fibers survived repeated bending, indicating a potential for robust surgical handling unlike what is typically seen for silicon, glass-encapsulated, or other metal penetrating electrodes [39]. Slight instrument adjustments in the surgical space often translated into large movements at the Flex Array–nerve interface, extreme enough for total carbon fiber breakage if not mitigated by a surgeon experienced with Flex Arrays. Biological motion surrounding peripheral nerve interfaces is likely more significant to intraneural electrode breakage than brain motion is for chronic penetrating brain electrodes. This array demonstrates a capability to withstand the long-term fatigue that a peripheral nerve interface is likely to experience by harnessin the intrinsic compliance of carbon fibers.

The blowtorch sharpening of carbon fibers resulted in a low-impedance electrode well within relevant electrophysiology recording range [45]. While SCFs had a 10.6 times larger surface area than laser cut carbon fibers (2735 μm^2^, 257 μm^2^, respectively) from Welle et al., 2020, blowtorching resulted in only a 1.7 times increase in 1 kHz impedance (32 kΩ, 19 kΩ, respectively). Despite the low 1 kHz impedance, it was unclear if the larger surface area would affect recording ability. We looked to similar devices to indicate the impact of a large surface area, such as the carbon nanotube yarn electrode. The yarn electrode recorded neural clusters from small autonomic nerves with electrodes exhibiting both a larger diameter (10–20 μm) and larger exposed length (200–500 μm), resulting in a larger surface area electrode [29], [33].

In fact, we recoded clear neural clusters with SCFs in rat CVN, feline DRG, and rat cortex. The amplitudes of neural clusters recorded from small, myelinated A fibers and unmyelinated C fibers within the rat CVN were predictably often lower than those recorded from cell bodies and axons within the feline DRG or rat cortex. The mean amplitudes of spontaneous neural clusters from the rat CVN, feline DRG, and rat cortex were 41.8 μV, 115.9 μV, and 80.7 μV, respectively. In addition to spontaneous activity, we recorded neural activity from the feline DRG corresponding to cutaneous brushing (152.4 ± 13.5 μV) and bladder pressure (86.8 ± 14.1 μV). The bladder clusters correlated to bladder pressure with a mean correlation coefficient of 0.61. This encouraging preliminary work, although not exhaustive, is the first example of carbon fiber electrode recordings in feline DRG, to our knowledge.

Novel, miniature, and implantable devices are required as bioelectric medicines advances in precise deciphering and monitoring of neural signals [1], [7]. The neural interface presented here could benefit a broad range of medical applications, such as control of hypertension [46], or monitoring of the digestive system [47] or cytokines [48]. Recording high quality, single units may improve the precision of these medical applications, especially with the CVN [49]. Due to their small diameter, carbon electrodes are more likely to land closer to regions of high electric field potential. This may lower stimulation thresholds below those used by other larger intraneural interfaces [50]. Promising preliminary stimulation results in other studies [37] enhance the prospect for this array in clinical peripheral nerve applications.

The array presented here uses stable, biocompatible materials appropriate for clinical use, such as carbon fiber [51] and medical-grade elastomer. Metal coatings traditionally used in medical devices, such as platinum iridium or iridium oxide [52], [53], could replace PEDOT:pTS to increase long-term stability. The large surface area of bare carbon electrode is favorable for neurotransmitter sensing [38], [54], [55]. With this larger surface area, we were able to record neural units. However, the smaller amplitude of neural clusters from the rat cortex recorded here as compared to our previous studies may be due to the larger surface area [36], [45]. A smaller surface area electrode might be achieved by electrochemical etching [56].

Additionally, this array’s archiecture may enable interfacing with difficult to access locations or small autonomic structures that require durable chronic interfaces [7]. The ability to insert individuated, sharpened fibers as an array without assistance may enable larger channel count arrays of small stiff electrodes [57], [58]. The ability to penetrate DRG suggests that these probes may be useful for other vertebrate ganglia, such as the nodose ganglia [59], and invertebrate ganglia, such as those in Aplysia [60]. The mechanical durability in a soft substrate makes these arrays ideal candidates for long-term studies in large animal models [61], [62]. Additionally, carbon fiber and carbon composite electrodes previously demonstrated a minimal chronic scarring response in brain [30], [38] and nerve [29].

There are, however, difficulties in deploying these devices for large-scale use. The manual fabrication of the carbon fiber arrays is a time-intensive process for an experienced fabrication technician. The application of silicone is limited by the small movements necessary to place the silicone in the wells and not on the fibers. The polyimide PCB and holder would likely require modifications to expand beyond four channels. While whole arrays can be sharpened at once – an improvement from the previous individual lasering technique [36] – the amount of exposed carbon can not yet be pre-determined. A chronic attachment method is still needed, such as suturing or Rose Bengal photochemical tissue bonding [63]. Future studies will need to investigate the chronic histological response of carbon fibers in peripheral structures.

## V. Conclusion

This work examines novel mechanical improvements to carbon fiber electrode arrays for interfacing with the nervous systems. SCFs can penetrate into stiff substrates, such as feline DRG and rat CVN, and reach deeper layers of the rat cortex, verifying their use with various recording targets despite their small diameter. The carbon fiber arrays embedded in silicone demonstrated durability beneficial for surgical handling and chronic peripheral nerve interfacing. Our electrophysiology recordings from the DRG were the first reported recordings with carbon fiber electrodes, to the best of our knowledge. The ease of insertion and durability to movement make these arrays strong candidates for future chronic *in vivo* use.

## Supporting information

Welle Supplemental 1

Welle Supplemental 2

Welle Supplemental 3

Welle Supplemental 4

## Acknowledgments

The authors thank Dr. Stephen Kemp for experimental guidance, Dr. Lauren Zimmerman for initial testing, Eric Kennedy for surgical preparation, David Ratze for experimental assistance, and Hao Shen and Hope Thayer for fabrication assistance. The authors acknowledge the financial support of the University of Michigan College of Engineering and NSF grants #DMR-0320740 and #DMR-1625671. The authors thank the staff at the Michigan Center for Materials Characterization and the Lurie Nanofabrication Facility for their technical support.

